# DIRECT RT-qPCR DETECTION OF SARS-CoV-2 RNA FROM PATIENT NASOPHARYNGEAL SWABS WITHOUT AN RNA EXTRACTION STEP

**DOI:** 10.1101/2020.03.20.001008

**Authors:** Emily A. Bruce, Meei-Li Huang, Garrett A. Perchetti, Scott Tighe, Pheobe Laaguiby, Jessica J. Hoffman, Diana L. Gerrard, Arun K. Nalla, Yulun Wei, Alexander L. Greninger, Sean A. Diehl, David J. Shirley, Debra G. B. Leonard, Christopher D. Huston, Beth D. Kirkpatrick, Julie A. Dragon, Jessica W. Crothers, Keith R. Jerome, Jason W. Botten

## Abstract

The ongoing COVID-19 pandemic has caused an unprecedented need for rapid diagnostic testing. The Centers for Disease Control and Prevention (CDC) and the World Health Organization (WHO) recommend a standard assay that includes an RNA extraction step from a nasopharyngeal (NP) swab followed by reverse transcription-quantitative polymerase chain reaction (RT-qPCR) to detect the purified SARS-CoV-2 RNA. The current global shortage of RNA extraction kits has caused a severe bottleneck to COVID-19 testing. We hypothesized that SARS-CoV-2 RNA could be detected from NP samples via a direct RT-qPCR assay that omits the RNA extraction step altogether, and tested this hypothesis on a series of blinded clinical samples. The direct RT-qPCR approach correctly identified 92% of NP samples (n = 155) demonstrated to be positive for SARS-CoV-2 RNA by traditional clinical diagnostic RT-qPCR that included an RNA extraction. Thus, direct RT-qPCR could be a front-line approach to identify the substantial majority of COVID-19 patients, reserving a repeat test with RNA extraction for those individuals with high suspicion of infection but an initial negative result. This strategy would drastically ease supply chokepoints of COVID-19 testing and should be applicable throughout the world.

## MAIN

The ongoing COVID-19 pandemic has put exceptional strain on public health laboratories, hospital laboratories, and commercial laboratories as they attempt to keep up with demands for SARS-CoV-2 testing. The current diagnostic testing methods recommended by the Centers for Disease Control and Prevention (CDC) in the United States and the World Health Organization (WHO) are traditional RT-qPCR assays that require two steps: first, an RNA extraction from patient nasopharyngeal (NP) swab material, followed by RT-qPCR amplification of the extracted RNA to detect viral RNA ^1-3^. The major bottleneck to widespread SARS-CoV-2 testing lies at the RNA extraction step. The simplest manual kit (the Qiagen Viral RNA Mini) is no longer available, and reagents and supplies for the larger automated instruments are extremely limited with uncertain supply chains. While substitution of other RNA extraction kits ^4,5^ is possible, they too are in limited supply. The current bottleneck is not simply the availability of RNA extraction kits, but also the cost of the extraction assay, the labor and time required to perform it, and the fact that it is rate limiting compared to the downstream RT-qPCR analysis. To address this issue, we tested the unconventional approach of skipping the RNA extraction step altogether and directly loading patient swab material into the RT-qPCR mix. Herein, we report that this approach (which we refer to hereafter as “direct RT-qPCR”) correctly identified 92% of samples (n =155) previously shown to be positive for SARS-CoV-2 RNA by conventional RT-qPCR featuring an RNA extraction. Thus, our results suggest that this streamlined assay could greatly alleviate constraints to COVID-19 testing in many regions of the world.

We initially conducted a pilot experiment using NP swabs from two COVID-19 patients who had previously been verified for SARS-CoV-2 infection by the Vermont Department of Health Laboratory (VDHL) using the CDC’s recommended RT-qPCR test. Both patient samples, which were originally collected as NP swabs in 3 mL of M6 viral transport medium (termed diluent hereafter), were pooled (equi-volume). RNA was extracted from 140 µL of the pooled sample using the QIAamp Viral RNA Mini kit, and purified RNA representing 11.3 µL of the original swab diluent was detected as positive via standard RT-qPCR using the CDC 2019-nCoV N3 primer/probe set, with a CT of 18.7. In parallel, we added 7 µL of the pooled COVID-19 patient NP swab diluent directly to the RT-qPCR reaction, and found that SARS-CoV-2 RNA was successfully detected in the absence of an RNA extraction step. Compared to the same pooled NP swab diluent extracted with the QIAamp Viral RNA Mini kit (after adjusting for the quantity of swab diluent added in each case), adding the NP diluent directly into the RT-qPCR reaction resulted in an ∼ 4 CT drop in sensitivity (**Fig. 1a**). Preheating the NP diluent for five minutes at 70°C prior to RT-qPCR had no impact on viral RNA detection. These results provided proof-of-principle that successful detection of SARS-CoV-2 RNA from an NP swab sample by RT-qPCR could be done in the absence of an RNA extraction step.

**Fig. 1.**
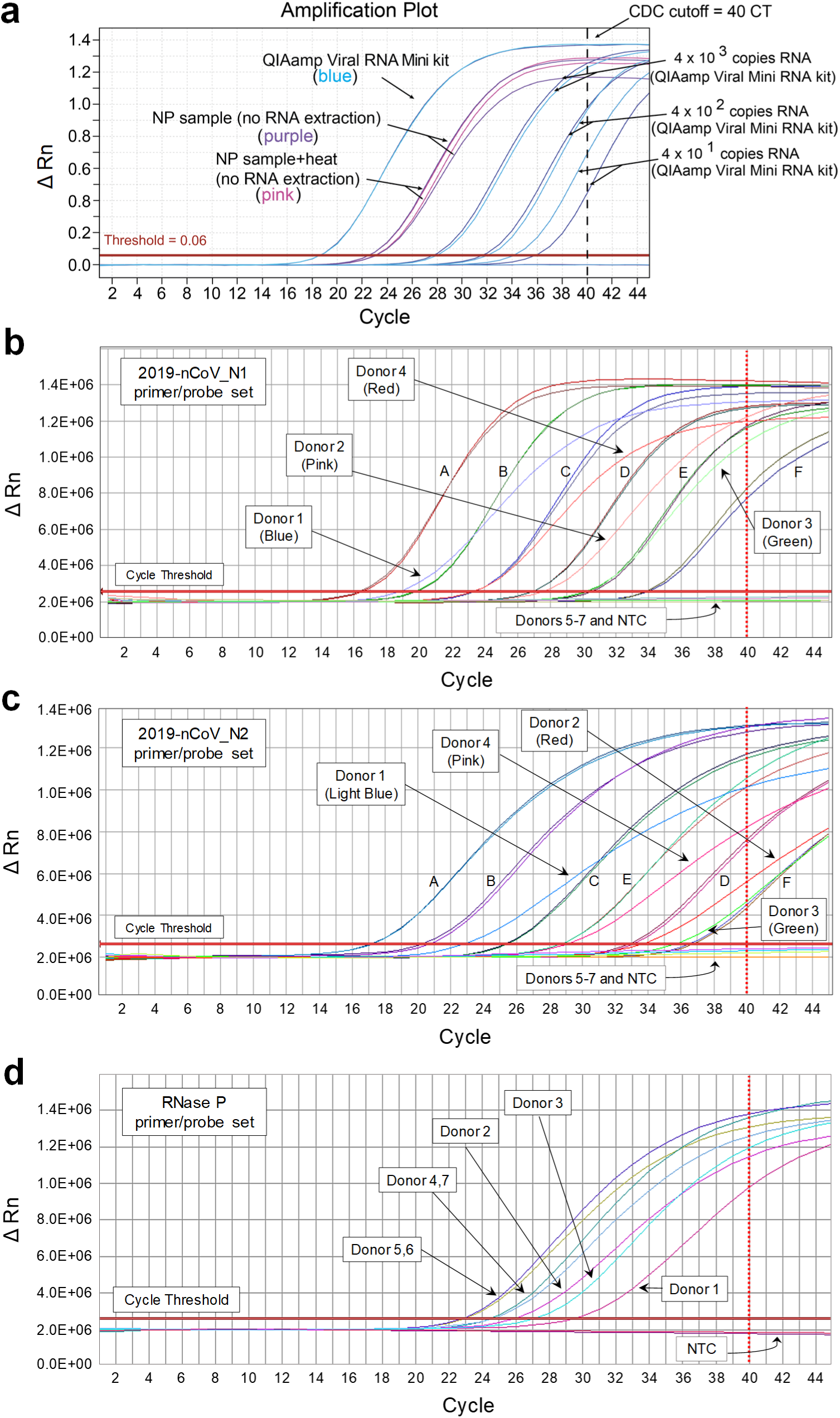
SARS-CoV-2 RNA can be detected from COVID-19 patient nasopharyngeal swabs by RT-qPCR without an RNA extraction step. (**a**) Nasopharyngeal (NP) swab diluents from two confirmed COVID-19 patients were pooled, and using the 2019-nCoV_N3 primer/probe set, the mixture was either i) subjected to RNA extraction using the Qiagen QIAamp Viral RNA Mini kit followed by subsequent testing by RT-qPCR (using the equivalent of 11.3 ul swab diluent) or ii) directly added to the RT-qPCR reaction, with or without a preheating step (five minutes at 70°C, “NP sample + heat”). As a control, the indicated quantities of the CDC 2019-nCoV Positive Control SARS-CoV-2 synthetic RNA was spiked into M6 transport media, purified using the QIAamp Viral RNA Mini kit, and screened by RT-qPCR. NP swab samples from seven additional donors were screened by direct RT-qPCR for SARS-CoV-2 RNA using (**b**) the 2019-nCoV_N1 primer/probe set, (**c**) the 2019-nCoV_N2 primer/probe set, or (**d**) for human RNase P RNA using the RP primer probe set. NP swab samples from donors 1 – 4 were previously shown to contain SARS-CoV-2 RNA by standard clinical RT-qPCR, while donors 5 – 7 were negative. For each primer/probe set, 7 µL (**a**) or 3 µL (**b, c, d**) of NP swab diluent was tested in the RT-qPCR reaction per donor. For the N1 and N2 primer probe sets, the fully synthetic SARS-CoV-2 RNA Control 2 from Twist Bioscience was loaded at serial 10-fold dilutions (A, 3×10^6^ copies; B, 3×10^5^ copies; C, 3×10^4^ copies; D, 3×10^3^ copies; E, 3×10^2^ copies; F, 3×10^1^ copies) as indicated in panels **b** and **c**. No template control (NTC) wells were included for each primer/probe set and each was negative. For panels **b** and **c**, the correlation coefficients (R^2^) of the standard curves were 0.999 and 0.995, respectively. The dashed line at cycle 40 in each graph indicates the limit of detection.

We next sought to validate the direct RT-qPCR approach on additional samples, determine the optimal volume of NP swab diluent for use in the direct RT-qPCR assay, and further address the potential impact of a prior heating step on assay sensitivity. NP samples from three COVID-19 patients who had previously been shown to be positive for SARS-CoV-2 RNA at high, intermediate, or low copy load by the Department of Laboratory Medicine at the University of Washington (UW) in Seattle were heated or not at 95°C for 10 minutes and then directly loaded into RT-qPCR reactions at 1, 3, or 5 µL volumes, or subjected to RNA extraction via the Roche MagNA Pure 96 platform prior to loading the equivalent of ∼20 µL of swab diluent into the RT-qPCR reaction. The main findings of this experiment, shown in **Table 1**, were that i) SARS-CoV-2 RNA could be detected in all three viral copy level samples at either input volume by direct RT-qPCR, provided they were heated first, and ii) addition of less NP diluent led to more sensitive detection of target RNA. Thus, heating appears important for subsequent detection of low viral copy samples, presumably by denaturing inhibitors of the RT and/or PCR enzymes present in the NP diluent. The best sensitivity for SARS-CoV-2 detection was achieved when 3 µL of swab diluent was used for direct RT-qPCR **(Table 1)**.

**Table 1.** Detection of SARS-CoV-2 RNA from NP swab diluent by direct RT-qPCR and the impact of heat and loading volume on assay sensitivity.

| Sample | Direct RT-qPCR (no RNA extraction)<sup>a</sup> |  |  |  |  |  | RT-qPCR<sup>b</sup><br>(+RNA extraction) |  |
| --- | --- | --- | --- | --- | --- | --- | --- | --- |
| | 5 μL swab diluent | | 3 μL swab diluent | | 1 μL swab diluent | | 20 μL diluent equivalent | |
|  | 95°C<sup>c</sup> | No heat | 95°C | No heat | 95°C | No heat | 95°C | No heat |
| **NP #1** | 24.0 | 26.5 | 20.7 | 21.2 | 20.9 | 20.8 | 16.8 | 15.8 |
| **NP #2** | 28.6 | 33.6 | 26.1 | 25.7 | 26.4 | 27.0 | 22.1 | 20.3 |
| **NP #3** | 38.2 | NEG<sup>e</sup> | 33.1 | 33.7 | 33.2 | 33.8 | 28.5 | 26.5 |
| **UTM<sup>d</sup>** | NEG | NEG | NEG | NEG | NEG | NEG | NEG | NEG |

<sup>a</sup>The indicated volumes of NP swab diluent were loaded directly into RT-qPCR featuring the 2019-nCoV\_N2 primer/probe set and the resulting CT values of each sample are shown.
<sup>b</sup>Standard RT-qPCR assay. The equivalent of 20 μL of RNA extracted from each NP swab sample was loaded into RT-qPCR featuring the 2019-nCoV\_N2 primer/probe set and the resulting CT values of each sample are shown.
<sup>c</sup>NP swab diluent was either heated for 10 min at 95°C or not prior to either direct or standard RT-qPCR.
<sup>d</sup>UTM, universal transport medium
<sup>e</sup>NEG, sample was negative after 40 cycles of qPCR

The CDC RT-qPCR method includes an internal control primer/probe set for detection of human RNase P. To ensure that direct RT-qPCR was able to detect the presence of this gene in NP swab diluent, and to test whether our approach was generalizable to other primer/probe sets, we screened seven donors (n = 4 COVID-19 positive; n = 3 COVID-19 negative) using the 2019-nCoV_N1, 2019-nCoV_N2, and RNase P primer/probe sets. Both the 2019-nCoV_N1 and 2019-nCoV_N2 primer/probe sets specifically detected SARS-CoV-2 RNA in each sample from COVID-19 patients, but not in samples from the negative donors (**Fig. 1b,c**). In addition, RNase P RNA was successfully detected in each donor by direct RT-qPCR (**Fig. 1d**). Collectively, these results confirmed i) the validity of our direct RT-qPCR approach with additional virus-specific primer/probe sets for detection of SARS-CoV-2 RNA, and ii) the presence of intact cellular RNA, as detected with an internal primer-probe set targeting a host gene, in all samples tested.

To get an accurate view of how omission of the RNA extraction would perform in a real-world clinical setting, we tested in a blinded fashion a panel of 150 NP swabs from COVID-19 patients representing high (CT less than 20), intermediate (CT of 20 - 30), or low (CT of more than 30) viral RNA loads by direct RT-qPCR. The SARS-CoV-2 RNA copy levels in these samples were previously determined at UW by standard clinical RT-qPCR that included RNA extraction. To address the potential impact of a prior heating step on detection sensitivity, NP swab samples were heated or not for 10 min at 95°C prior to use in the downstream RNA extraction and/or direct RT-qPCR. For each NP swab sample, a 3 µL volume of diluent was used for direct RT-qPCR. In parallel, a 30 µL aliquot of each sample was used for RNA extraction and an equivalent of 3 µL of the original swab diluent used for RT-qPCR, allowing for a one to one comparison with the direct RT-qPCR method. To control for inhibitors of the RT or PCR enzymes and/or RNase activity in the swab diluent, an aliquot of each swab diluent was spiked with 4 x 10^4^ copies of EXO control RNA for subsequent detection with an EXO primer-probe set; no inhibition was noted under any of the tested conditions. **Supplementary Table 1** shows the full results of this experiment while **Table 2** provides a summary. **Fig. 2** shows the CT values from the original clinical RT-qPCR done on the equivalent of 20 µL of NP swab diluent versus those obtained by direct or standard RT-qPCR when measuring 3 µL (or equivalent) of swab diluent. There were several key observations from this experiment. First, preheating NP swab samples prior to direct RT-qPCR enhanced assay sensitivity (e.g. 138 of 150 samples were detected following a heat step, compared with only 126 of 150 positive in the absence of preheating). Second, extraction of RNA prior to RT-qPCR did not enhance detection of SARS-CoV-2 positive samples when an equivalent amount of diluent was screened by direct RT-qPCR (138 of 150 samples were detected by each approach). This suggests one of the primary advantages of kit-based RNA extraction comes from concentrating the sample material so more RNA can be loaded into the RT-qPCR reaction. Third, 11 of the 12 samples not detected by direct RT-qPCR were from donors that had extremely low loads of viral RNA as originally determined in the clinical test (CTs ranging from 33 to 38). Fourth, preheating samples reduced sensitivity when using the Roche MagnaPure 96 RNA extraction system, but had little effect on the direct RT-qPCR approach. Finally, the results from the EXO spiking experiment indicate that the NP swab diluent does not inhibit the direct RT-qPCR assay. Collectively, these results demonstrate that the direct RT-qPCR assay described here is capable of reliably detecting the substantial majority of COVID-19 patients.

**Table 2.** Detection sensitivity of direct RT-qPCR versus standard RT-qPCR on NP swabs containing a range of SARS-CoV-2 viral RNA loads.

| Viral RNA load<sup>a</sup> | Direct RT-PCR (3 μL swab diluent)<sup>b</sup> | | Standard RT-qPCR (3 μL swab diluent equivalent)<sup>c</sup> | | Standard RT-qPCR (20 μL swab diluent equivalent)<sup>d</sup> | |
| --- | --- | --- | --- | --- | --- | --- |
|  | 95°C<sup>e</sup> | No heat | 95°C | No heat | 95°C | No heat |
| **High (CT < 20)** | 30/30<br>(100%) | 30/30<br>(100%) | 30/30<br>(100%) | 30/30<br>(100%) | 16/16<br>(100%) | 16/16<br>(100%) |
| **Intermediate (CT 20 – 30)** | 102/103<br>(99%) | 94/103<br>(91%) | 102/103<br>(99%) | 103/103<br>(100%) | 74/74<br>(100%) | 74/74<br>(100%) |
| **Low (CT > 30)** | 6/17<br>(35%) | 2/17<br>(12%) | 6/17<br>(35%) | 10/17<br>(59%) | 6/10<br>(60%) | 8/10<br>(80%) |
| **Total** | **138/150**<br>**(92%)** | **126/150**<br>**(84%)** | **138/150**<br>**(92%)** | **143/150**<br>**(95%)** | **96/100**<br>**(96%)** | **98/100**<br>**(98%)** |

<sup>a</sup>CTs determined by clinical RT-qPCR at UW Seattle using the equivalent of 20 μL of RNA extracted from an NP swab. The 2019-nCoV\_N2 primer/probe set was used for the RT-qPCR reactions.
<sup>b</sup>The indicated volume of NP swab diluent was loaded directly into RT-qPCR featuring the 2019-nCoV\_N2 primer/probe set
<sup>c</sup>RNA was extracted from 30 μL of NP swab diluent and the equivalent of 3 μL NP swab diluent was loaded into RT-qPCR featuring the 2019-nCoV\_N2 primer/probe set.
<sup>d</sup>RNA was extracted from 200 μL of NP swab diluent and the equivalent of 20 μL NP swab diluent was loaded into RT-qPCR featuring the 2019-nCoV\_N2 primer/probe set.
<sup>e</sup>NP swab diluent was either heated for 10 min at 95°C or not prior to either direct RT-qPCR or RNA extraction followed by standard RT-qPCR.

**Fig. 2.**
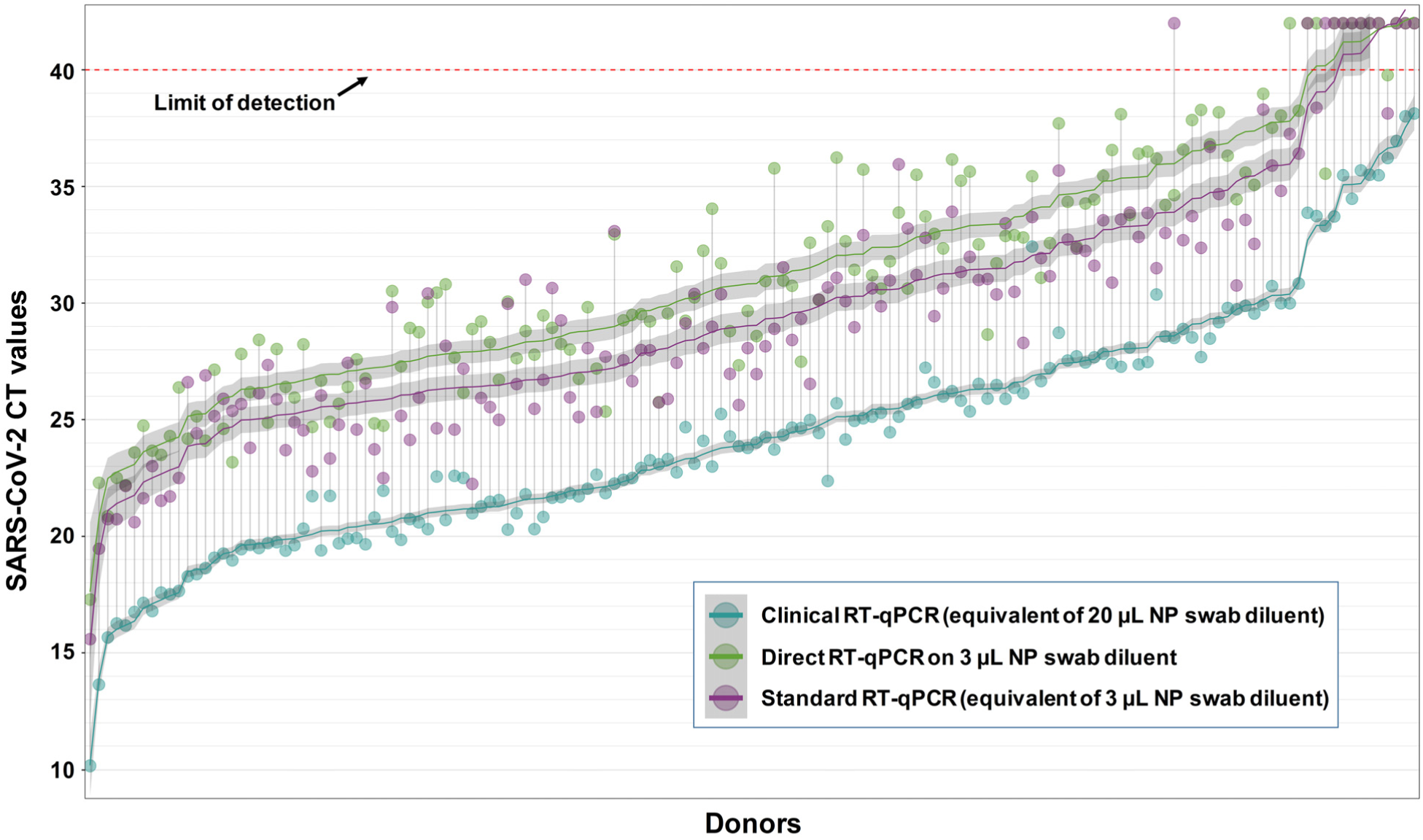
Distribution of CT values from COVID-19 patient NP swabs following direct RT-qPCR versus standard RT-qPCR that includes RNA extraction. 150 NP swab samples representing high (CT values less than 20), intermediate (CT values of 20-30), or low (CT values of more than 30) SARS-CoV-2 RNA loads as determined by standard clinical RT-qPCR at UW in Seattle (aqua circles) were analyzed by the indicated method. All assays used the 2019-nCoV_N2 primer/probe set. Direct RT-qPCR was performed on 3 µL of NP swab diluent after heating for 10 minutes at 95°C (green circles). In parallel, RNA was extracted from 30 µL of NP swab diluent that had been previously heated at 95°C for 10 minutes, and RNA representing 3 µL of the original diluent was used in RT-qPCR (purple circles) to allow a head to head comparison to direct RT-qPCR on the same quantity of NP swab diluent. The limit of detection (40 CT) is denoted with a dashed line. Samples with CT values above this cutoff were considered negative for SARS-CoV-2 RNA. The fitted curves are LOESS-smoothed (locally estimated scatterplot smoothing) CT values, with 95% confidence interval in gray, against the mean of CT values detected in the clinical RT-qPCR assay with primer sets N1 & N2. Samples are ordered by the latter mean. The full data set for this experiment and controls is provided in **Supplementary Table 1**.

COVID-19 testing demands are currently overwhelming the world’s clinical laboratories. A major chokepoint is the RNA extraction step, due to both the time and labor for this step and the critical shortage of extraction kits and required reagents. As a means to circumvent these limitations on clinical testing availability, we have shown that a direct RT-qPCR approach that omits an RNA extraction step can effectively identify SARS-CoV-2 RNA from NP swabs. Further, direct RT-qPCR is able to detect host RNA from NP swabs, which enables us to rule out false negatives due to inadequate swabbing of patients resulting in a paucity of intact RNA in the specimen, or due to the presence of inhibitors. When applied to a large collection of clinical NP specimens representative of the range of COVID-19 patients in the state of Washington, the direct RT-qPCR assay correctly identified 92% of the donors screened. The only donors that our simplified approach missed were those with the very lowest load of viral RNA (and who were on the very cusp of detection even when RNA extraction was used). Of the 1,872 cases with a positive diagnosis at UW by our team at the time of this writing, only ∼25% would fall into this low copy range, which indicates that this assay will detect the substantial majority of COVID-19 patients. Indeed, after we posted our initial results on bioRxiv ^4^, two additional groups posted preliminary data on bioRxiv and medRxiv replicating these findings, using standard clinical diagnostic methods in Chile and Denmark ^6,7^. The simplified approach we present here may be especially well suited for general screening of the public and identification of “silent carrier” individuals. Where resources permit, we envision that a two-step testing algorithm in which a repeat test including RNA extraction could be applied for symptomatic patients, health care workers, and others with a high index of suspicion but an initial negative result. Such a two-step strategy would reduce the need for RNA extraction for a substantial portion of future COVID-19 tests. Importantly, many testing sites in the developing world may not even have the capacity to perform RNA extraction, but could easily run RT-qPCR reactions. This new approach would open up a viable avenue for testing in these regions.

## Methods

### Samples

#### University of Vermont

Clinical nasopharyngeal swabs were collected in 3 ml M6 transport medium. Patients were confirmed to be negative (3) or positive (6) for COVID-19 by the Vermont Department of Health using the CDC 2019-Novel Coronavirus (2019-nCoV) Real-Time RT-PCR Diagnostic Panel ^1^. Limited quantities of this material were available, so we initially pooled equal amounts of sample from two COVID-19 patients (**Fig. 1a**). Subsequently, four additional COVID-19 confirmed patients and four COVID-19 negative donors were tested individually for the presence of SARS-CoV-2 RNA or RNase P by direct RT-qPCR of the sample material (**Figs. 1b-d**).

#### University of Washington

Clinical nasopharyngeal swabs were collected in 3 mL of Universal transport medium (UTM). 150 samples from patients confirmed to be COVID-19 positive by the University of Washington Medical Laboratory with ranges of viral loads (CT values of SARS-CoV-2 with the N2 primer from the original clinical test run at UW range from 10.17 to 38.13) and sufficient volume were selected for this study (**Fig. 2, Tables 1 and 2, and Supplementary Table 1**).

All patient samples at each testing site were stored at 4 oC in between sample collection and transport to the laboratory, and again until clinical testing was carried out, maintaining a standardized sample acquisition and processing protocol.

### RNA Extraction

#### University of Vermont

To compare the effect of nucleic acid extraction to direct RT-PCR, RNA from a pooled patient sample was extracted using the QIAamp Viral RNA Mini (Qiagen, Cat. No. 52904) kit according to the manufacturer’s instructions. 140 µL of pooled nasopharyngeal swab diluent was extracted and eluted in 60 µL; 5 µL (equivalent to 11 µL of original sample) was used for real-time RT-PCR (**Fig. 1a**).

#### University of Washington

To compare the effect of nucleic acid extraction, RNA from 30 µL or 200 µL of patient sample was extracted using the Roche MagNA Pure 96 (Roche Lifesciences) platform, eluted into 50 µL of buffer; 5 µL of RNA (equivalent to 3 µL of original sample for a 30 µL extraction) was used for real-time RT-PCR. To monitor RNA recovery and RT-PCR efficiency, 40,000 copies of EXO internal control RNA were added into the lysis buffer and went through the extraction process with each sample (**Fig. 2, Tables 1 and 2, and Supplementary Table 1**).

### RT-qPCR Detection

#### University of Vermont

In **Fig. 1a**, 7 µL of pooled nasopharyngeal swab diluent (heated to 70°C for 5 minutes, or not), or 5 µL extracted RNA, was used as input material for the New England Biolabs Luna Universal Probe One Step RT qPCR kit (Cat #E3006S, lot #10066679) according to the IDT recommendation for primers/probe (1.5 µL primer/probe per reaction) using primer set N3 from IDT’s 2019-nCoV CDC Emergency Use Authorization Kits (20 µL reaction). Subsequently, in **Figs. 1b-d**, 3 µL of nasopharyngeal swab diluent was used as input material for the ThermoFisher TaqPath™ 1-Step RT-qPCR Master Mix, CG (Cat #A15299) using primer sets N1, N2, and RP (human RNase P) from IDT’s CDC Emergency Use Authorization Kits (20 µL reaction). Known copies of positive control RNA (CDC 2019-nCoV Positive Control *in* vitro transcribed RNA for **Fig. 1a**; fully synthetic SARS-CoV-2 RNA Control 2 [Cat# MN908947.3, Twist Bioscience] for **Fig. 1b,c**) were used to compare viral RNA quantities in patient samples. Each patient sample, as well as no template control (water as well as M6 transport media) was run in duplicate, the synthetic RNA standard was run in triplicate. The initial experiment was run on an ABI QuantStudio 6 Flex (**Fig. 1a**), while subsequent experiments were run on an ABI 7500 Fast (**Figs. 1b-d**). Reagent preparations were performed in a PCR-free clean room equipped with PCR workstations (AirClean Systems). Clinical sample manipulations (RNA extractions or direct addition of NP swab diluent to RT-qPCR plates) were done in a Class IIA Biosafety Cabinet using BSL-2 precautions. RT-qPCR plates were placed directly in biosafety waste at the conclusion of each run.

#### University of Washington

**Fig. 2, Tables 1 and 2, and Supplementary Table 1**: For each patient sample, 250 µL was heat treated at 95oC for 10 minutes, or not. For direct RT-PCR, 3 µL of this sample, or 5 µL of extracted RNA, was added directly into the RT-PCR reaction mixture. Each real-time RT-PCR reaction contained 400 nM of CDC N2 forward and reverse primers and 100 nM of N2 probe. To monitor potential RT-PCR inhibition, each RT-PCR reaction was spiked with 40,000 copies of EXO internal control RNA and EXO primers (100 nM of EXO forward and 200 nM of EXO reverse) and probes (62.5 nM). Real-time RT-PCR assays were performed using AgPath-ID One Step RT-PCR kit (Life Technologies) and an ABI 7500 Real Time PCR system was used to perform the RT-PCR reactions ^8^. For samples that were subjected to RNA extraction, 40,000 copies of EXO RNA was spiked into the lysis buffer. Extraction efficiency for each sample was tracked based on the percentage of the spiked EXO RNA that was detected by RT-qPCR. The primer-probe sequences and RT-qPCR conditions for all methods used are below:

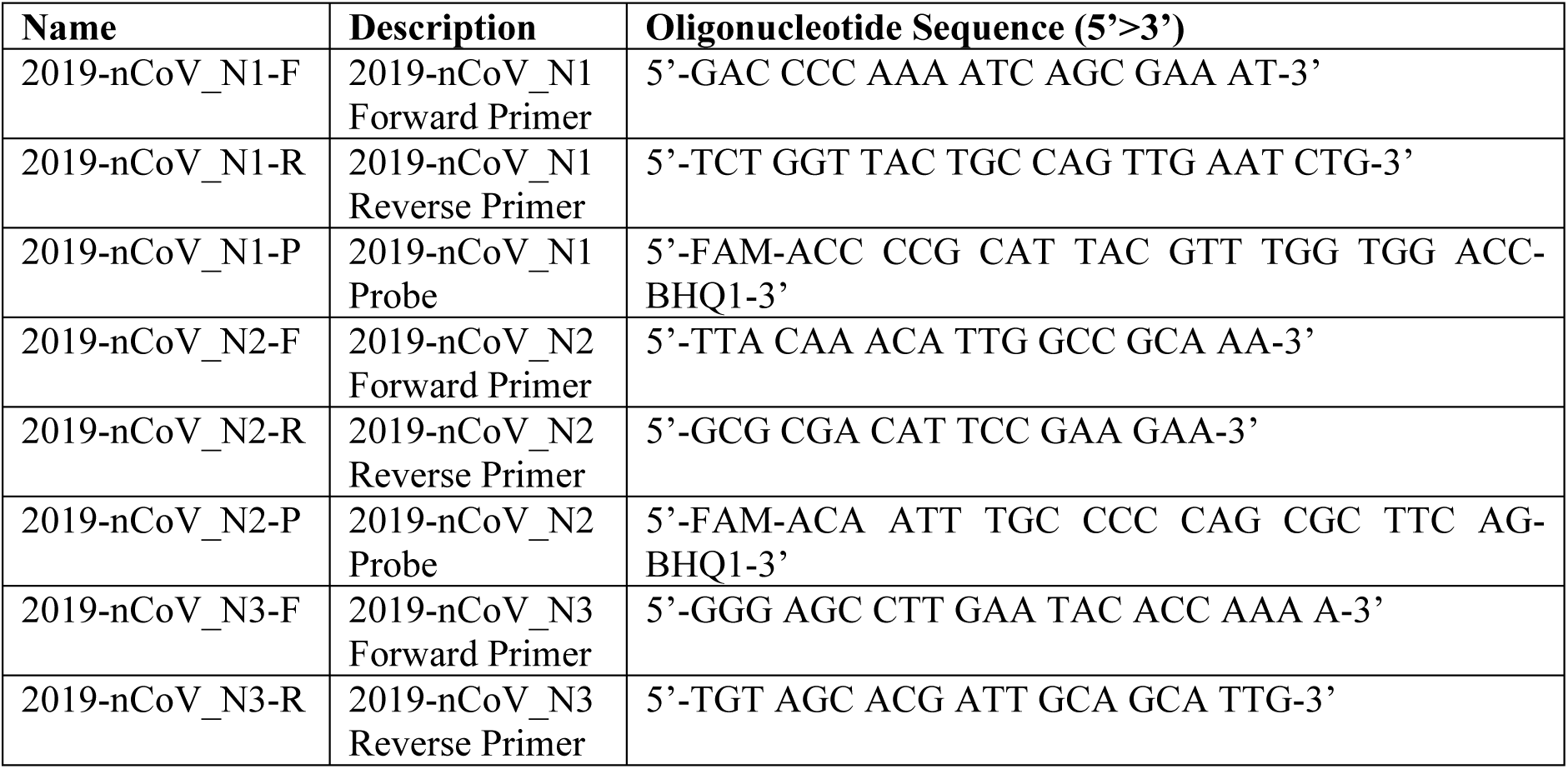
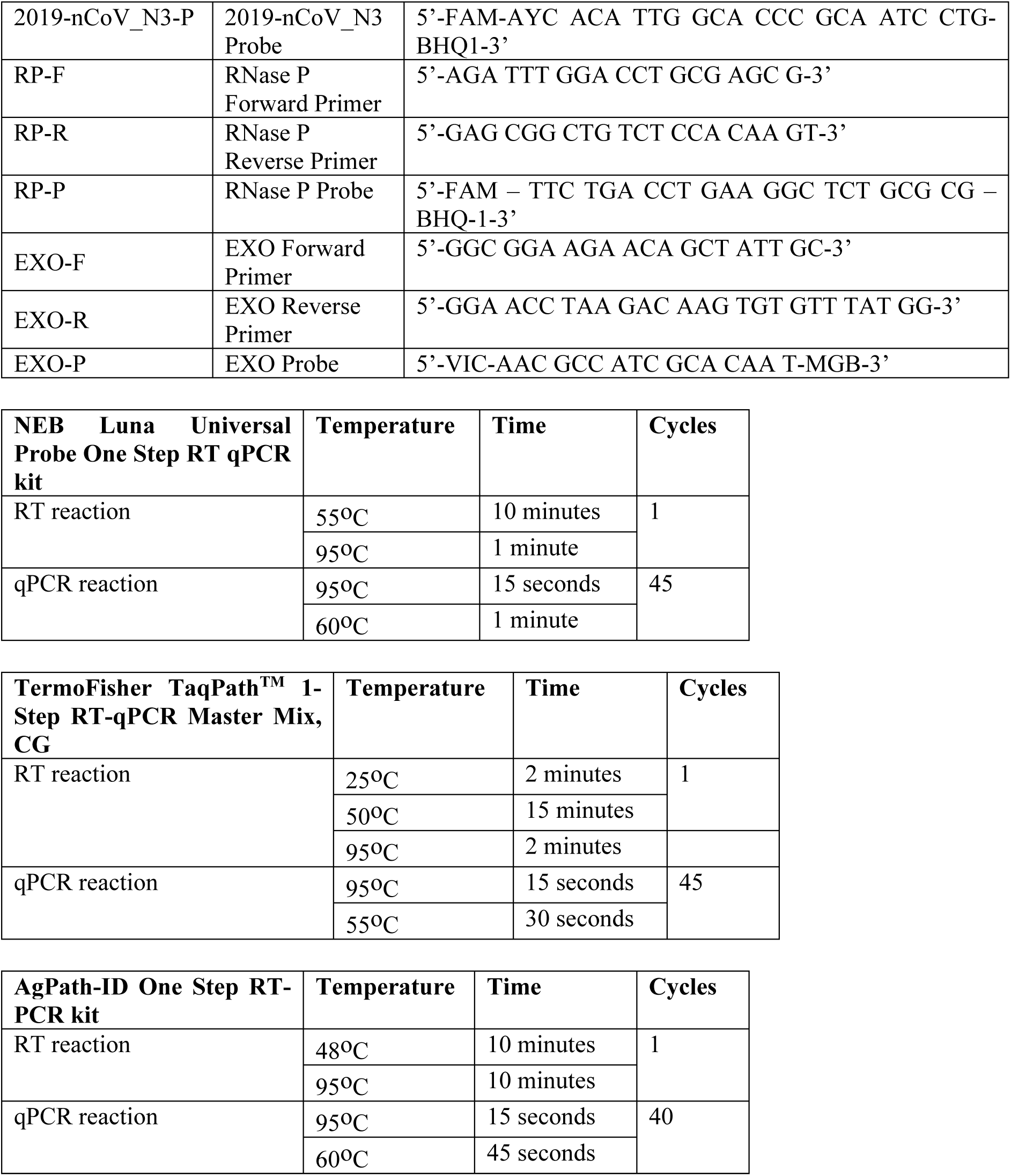

### Study Design

For both the University of Vermont and the University of Washington, sample selection (including information regarding CT values) and RNA extraction/RT-PCR of samples were performed by separate individuals; the person running the assays was blinded to the original clinical CT value of the samples.

## Acknowledgements

The authors gratefully acknowledge funding from the Office of the Vice President for Research at the University of Vermont (JWB), the M.J. Murdock Charitable Trust (KRJ), and NIH grants P30GM118228-04 (Immunology and Infectious Disease COBRE) (EB, JAD), P20GM125498-02 (Translational Global Infectious Diseases Research COBRE) (BDK, CDH, SAD, JAD), U01AI1141997 (JWB, SAD, and BDK), R41AI132047 (JWB), and U54 GM115516-01 (JAD). We also thank the staff at the Vermont Department of Health involved in COVID-19 diagnostic testing in particular Helen Reid, Cheryl Achilles, and Christine Matusevich as well as Dr. Deborah Donnell, Dr. Ken Hampel, Dr. Colm Atkins, Dr. Kishana Taylor, Dr. Robyn Kent, Daniel Vellone, and Denise Francis for technical comments or experimental help.

## Author Contributions

EAB, MLH, ST, KRJ, and JWB conceived and designed experiments. EAB, MLH, GAP, YW, ST, PL, JH, DG performed experiments. JWC, ALG, and KRJ provided specimens. EAB, ST, DJS, MLH, KRJ, and JWB analyzed the data. EAB, CDH, KRJ, and JWB wrote the manuscript and all authors participated in the editorial process and approved the manuscript.

## Ethical Statement

Patient samples were de-identified and were not considered Human Subjects Research due to the quality improvement and public health intent of the work.

